# Hedgehogs are the major amplifying hosts of severe fever with thrombocytopenia syndrome virus

**DOI:** 10.1101/2022.01.30.478379

**Authors:** Chaoyue Zhao, Xing Zhang, Junfeng Hao, Ling Ye, Kevin Lawrence, Yajun Lu, Chunhong Du, Xiaoxi Si, Haidong Xu, Qian Yang, Qianfeng Xia, Guoxiang Yu, Fei Yuan, Jiafu Jiang, Aihua Zheng

**Affiliations:** State Key Laboratory of Integrated Management of Pest Insects and Rodents, Institute of Zoology, Chinese Academy of Sciences, Beijing, 100101, China; CAS Center for Excellence in Biotic Interactions, University of Chinese Academy of Sciences, Beijing, 100049, China; College of life sciences, University of Chinese Academy of Sciences, Beijing, 100049, China; Core Facility for Protein Research, Institute of Biophysics, Chinese Academy of Science, Beijing, 100101, China; Daishan Center for Disease Control and Prevention, Zhoushan, Zhejiang, 316200, China; School of Veterinary Science, Massey University, Palmerston North 4442, New Zealand; Key Laboratory of Tropical Translational Medicine of Ministry of Education, School of Tropical Medicine and Laboratory Medicine, Hainan Medical University, Haikou, Hainan, 571199, China; Yunnan Institute for Endemic Diseases Control and Prevention, Dali, Yunnan, 671000, China; College of life sciences, Henan Normal University, Xinxiang, Henan, 453007, China; Shaozhuang central middle school, Qingzhou, Weifang, Shandong, 262507, China; Department of Infectious Disease, Yidu Central Hospital of Weifang, Weifang, Shandong, 252550, China; Changdao National Nature Reserve Management Center, Yantai, Shandong, 234000, China; State Key Laboratory of Pathogen and Biosecurity, Beijing Institute of Microbiology and Epidemiology, Academy of Military Medical Sciences, Beijing, 100071, China

**Keywords:** SFTSV, hedgehog, *Haemaphysalis longicornis*, transmission, host

## Abstract

Severe fever with thrombocytopenia syndrome virus (SFTSV) is a tick-borne bandavirus mainly transmitted by *Haemaphysalis longicornis* in East Asia, mostly in rural areas. To date, the amplifying host involved in the natural transmission of SFTSV remains unidentified. Our epidemiological field survey conducted in endemic areas in China showed that hedgehogs were widely distributed, had heavy tick infestations, and had high SFTSV seroprevalence and RNA prevalence. After experimental infection of *Erinaceus amurensis* and *Atelerix albiventris* hedgehogs with SFTSV, robust but transitory viremias were detected, which lasted for around nine to eleven days. The infected hedgehogs experienced light weight loss and histopathology of the spleen showed hemorrhagic necrosis and lymphopenia, with infected hedgehogs recovering after viral clearance. Remarkably, SFTSV transmission cycle between hedgehogs and nymph/adult *H. longicornis* was easily accomplished under laboratory condition with 100% efficiency. Furthermore, naïve *H. longicornis* ticks could be infected by SFTSV-positive ticks co-feeding on naïve hedgehogs, with transstadial transmission of SFTSV also confirmed. We also found that SFTSV viremia remained high in hedgehogs during hibernation, suggesting that this mechanism might contribute to the persistence of SFTSV from one year to the next. Of concern, we recently found evidence of the natural circulation of SFTSV in the urban area of Beijing City in China involving *H. longicornis* ticks and *E. amurensis* hedgehogs. Our study suggests that the hedgehogs are the major wildlife amplifying hosts of SFTSV and that urban outbreaks of SFTSV might occur in the future.

## Introduction

Severe fever with thrombocytopenia syndrome virus (SFTSV) is a new tick-borne bandavirus first identified in China in 2009(Yu et al., 2011), followed by Korea in 2011(Denic et al., 2011), Japan in 2014(Takahashi et al., 2014), Vietnam in 2019(Tran et al., 2019) and Pakistan in 2020(Zohaib et al., 2020). The symptoms of SFTS include fever, thrombocytopenia, leukocytopenia, and gastrointestinal disorders, with a case-fatality rate of between 2 and 30%(Liu, He, Huang, Wei, & Zhu, 2014; S. Liu et al., 2014; Yu et al., 2011). The earliest Chinese cases were reported in the Dabie mountain range, which is located at the intersection of Henan, Hubei, and Anhui provinces in central China. Besides the Dabie mountain range, Shandong, Liaoning, and Zhejiang provinces are the other main hot spots for SFTS in China(J. Sun et al., 2018). Within Zhejiang Province, Daishan County, an archipelago of islands located in the East China Sea, is one of the most endemic areas(Fu et al., 2016). The major industry in Daishan County is fishing and tourism, agriculture is relatively unimportant with only 4000 sheep and 150 cattle on the islands in 2019, as reported by the local government. As of 2020, SFTS cases have been reported in most other Chinese provinces(Lin et al., 2020; J. Sun et al., 2018; Zhu et al., 2019).

*Haemaphysalis longicornis* (Asian long-horned tick) is the major vector for SFTSV and the dominant human-biting tick in the SFTSV endemic areas(Li et al., 2016; Yun et al., 2015; G. Zhang, Zheng, Tian, & Li, 2019). *H. longicornis* has both bisexual and parthenogenetic populations, with the parthenogenetic populations being widely distributed in China and strongly correlated with the distribution of SFTS cases (X. Zhang et al., 2022). *H. longicornis* ticks go through a three-stage life cycle (larva, nymph, and adult). At each stage, they feed on a wide range of wild and domestic animals including mammals, birds, companion animals and livestock(Zhao et al., 2020).

Extensive reports suggest that *H. longicornis* is the reservoir of SFTSV(Luo et al., 2015; S. W. Park et al., 2014; Zhuang et al., 2018). However, the transstadial transmission efficiencies of SFTSV from egg pools to larvae pools, larval pools to nymph pools and nymph pools to adults were 80%, 92%, 40% or 100%, 100%, 50% under laboratory conditions according to two reports(Y. Y. Hu et al., 2020; Zhuang et al., 2018). Correspondingly, the SFTSV prevalence was extremely low in different developmental stages of host-seeking *H. longicornis* ticks collected from vegetation, ranging from 0.2% to 2.2%(Luo et al., 2015; S. W. Park et al., 2014; Wang et al., 2015). These findings suggest that ticks alone are not sufficient to maintain a reservoir of SFTSV in the natural environment, therefore one or more additional amplifying hosts are required.

Antibodies to SFTSV and viral RNA have been detected in a wide range of domestic animals, including goats, cattle, dogs, and pigs, and wild animals such as shrews, rodents, weasels, and hedgehogs. The highest seroprevalence was found in sheep (69.5%), cattle (60.4%), dogs (37.9%) and chickens (47.4%)(Chen et al., 2019; Huang et al., 2019; Niu et al., 2013). Given that most of the SFTS patients are farmers, who have frequent contacts with many of the domestic and wild animals listed above, this makes understanding the epidemiology of SFTSV both difficult and complex.

Hedgehogs belong to the family *Erinaceinae*, which includes twenty four genera and are widely distributed in the Eurasian continent and Africa(He et al., 2012). Some genera have even been introduced into countries with no indigenous hedgehogs, including Japan and New Zealand(Brockie, 1975; ISAAC, 2005). The Amur hedgehog *Erinaceus amurensis* is closely related to the European hedgehog, *Erinaceus europaeus*, but is slightly bigger and lighter in color. It is native to Amur Oblast and Primorye in Russia, the Korean Peninsula, and is common in northern and central China. The African pygmy hedgehog *Atelerix albiventris*, native to central and eastern Africa, has been introduced into many countries as pets, including China, where they are available in many suburban petshops(Brockie, 1975; ISAAC, 2005). Both the Amur hedgehog and the African pygmy hedgehog can become heavily infested by all kinds of ticks and are known to carry many zoonotic diseases, such as Tickborne encephalitis virus, Bhanja virus, and Tahyna virus(Dziemian, Sikora, Pilacinska, Michalik, & Zwolak, 2015; Jahfari et al., 2017; Riley & Chomel, 2005). Hedgehogs are poikilothermal animals and hibernate during winter. During hibernation, their metabolism and immune system are suppressed (Bouma, Carey, & Kroese, 2010) which has led to the suspicion that hibernating hedgehogs contribute to the long-term persistence of these viruses(Simkova, 1966). A few previous studies reported that SFTSV antibodies and RNA were detected in *E. amurensis* in Shandong and Jiangsu Province. However, the prevalence of SFTV infection appeared low as compared to that in other animals such as goats, sheep, and cattle(Li et al., 2016; Y. Sun et al., 2017).

In China, the density of large wild animals is extremely low, especially in East China where SFTS is endemic. Instead, the most abundant wildlife in these areas are rodents and insectivores(Jiang, Liu, Wu, Jiang, & Zhou, 2017). However, the potential role of rodents in the transmission of SFTSV was refuted when it was shown that immunocompetent rodents cannot develop SFTSV viremia after artificial inoculation(Matsuno et al., 2017). In contrast, hedgehogs are the only small wild animals which consistently show high SFTSV seroprevalence, high density, as well as high *H. longicornis* infestation in the SFTS endemic areas(Li et al., 2014; Y. Sun et al., 2017). They are widely distributed both in the urban and wild ecosystem(Smith et al., 2010), which has led us to speculate that hedgehogs might play an important role in the natural circulation of SFTSV in China. To test this hypothesis, we first carried out an epidemiological survey to confirm the role of hedgehogs as potential wild amplifying hosts for SFTSV. Then a series of linked laboratory experiments were performed to investigate the susceptibility and tolerance of hedgehogs to SFTSV infection, and to establish the transmission of infection between hedgehogs and *H. longicornis* ticks at both the nymph and adult stages.

## Results

### Field survey of hedgehogs in SFTS endemic areas

To confirm the role of hedgehogs as potential wild amplifying hosts for SFTVS, we firstly performed an animal survey in Daishan County, an archipelago of islands in the East China Sea (Figure 1A). Daishan County is the worst affected area for SFTS in Zhejiang Province (Fu et al., 2016) and between 2011 and 2019 one hundred and thirty-three SFTS cases were reported by Daishan CDC. SFTS cases were reported on all the major Daishan County islands including Daishan Island, Qushan Island and Changtu Island, with the exception of Xiushan Island, even though Xiushan Island has a similar landscape, vegetation, and population density as Daishan Island, Qushan Island and Changtu Island (Fig. 1B). Small mammal traps were set, and the following numbers of wildlife caught, 33 on Daishan Island and 75 on Xiushan Island. On Daishan Island, 28% of the captured small mammals were *E. amurensis* (Amur hedgehog), 18% were *Rattus. Norvegicus* (brown rat), 36% were *Sorex araneus* (common shrew), and 18% were *Apodemus agrarius* (striped field mouse). On Xiushan Island no hedgehogs were caught, 48% of the small mammals caught were *R. norvegicus*, 44% were *S. araneus* and 8% were *Rattus losea* (lesser ricefield rat) (Fig. 1C). Antibody testing showed that 3/9 (33%) of *E. amurensis* hedgehogs from Daishan Island were positive for SFTSV (Fig. 1D). Hedgehogs are abundant in the two villages in Daishan Island, with an estimated population density of greater than 80 individuals per square kilometer based on the results of the trapping study (Table 1). In addition, these nine trapped hedgehogs were all heavily infected by ticks with an average of 145 ticks per hedgehog, including *H. longicornis* (Table 2).

**Table 1.**
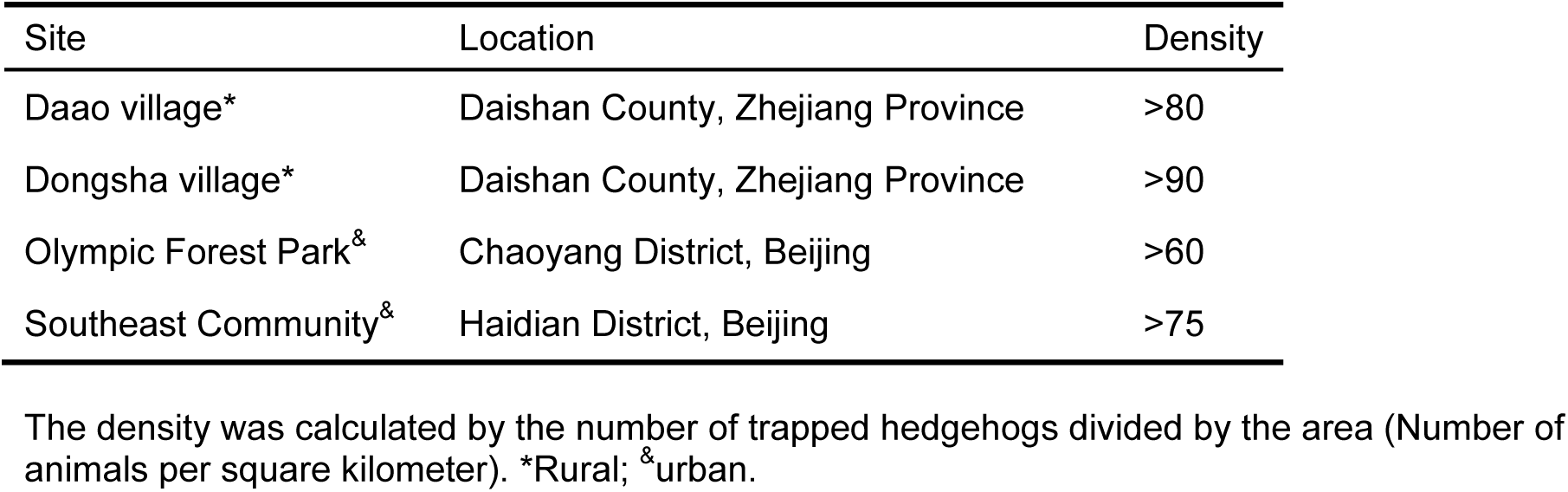
The density of hedgehogs in rural and urban areas.

**Table 2.** Average number of ticks collected from wild mammals captured in Daishan County.

| Animal | Number |
| --- | --- |
| *Sorex araneus Linnaeus* | 1.5 |
| *Rattus norvegicus* | 1 |
| *Rattus losea* | 0 |
| *Apodemus agrarius* | 0 |
| *Erinaceus amurensis* | 145 |

**Figure 1.**
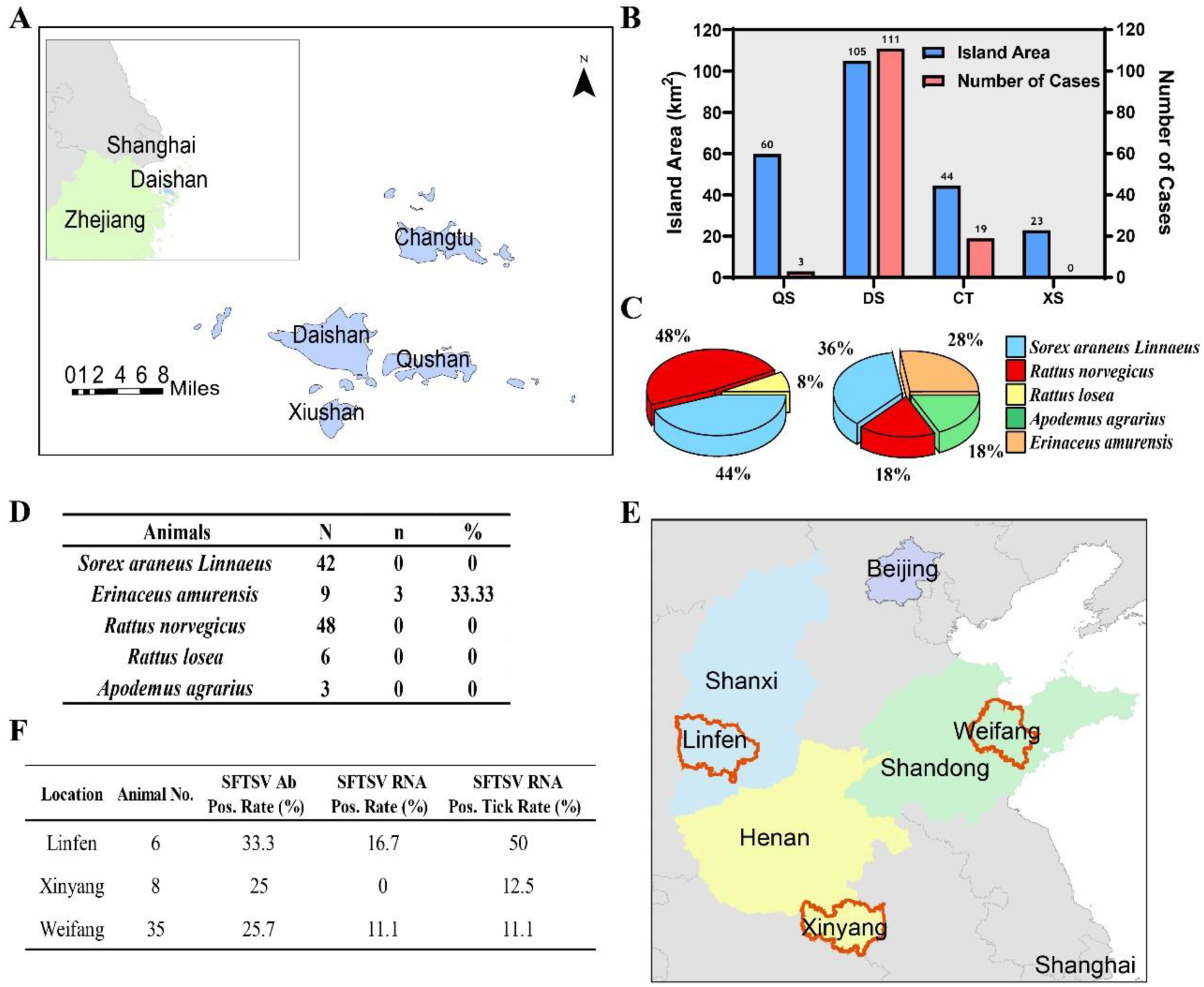
The association between hedgehogs and SFTSV endemic. (A) Locations of the major islands of Daishan County, Zhejiang Province. (B) Land area and SFTS case numbers for major islands in Daishan County. DS, Daishan Island; QS, Qushan Island; XS, Xiushan Island; CT, Changtu Island. (C) Species and relative rate of wild animals collected on Xiushan Island (left) and Daishan Island (right). (D) Seroprevalence of SFTSV in wild animals captured in Xiushan Island and Daishan Island. N, number of sampled animals; n, number of sampled animals positive for SFTSV antibody; %, percentage of sampled animals positive for SFTSV antibody. (E) Locations of Weifang City of Shandong Province, Linfen City of Shanxi Province, Xinyang City of Henan Province, where hedgehogs were collected. (F) Epidemiological analysis of trapped animals. Seroprevalence of SFTSV in hedgehogs was measured by ELISA against SFTSV nucleocapsid protein. Viral RNA was tested by PCR. Pos. is the abbreviation of positive. “SFTSV RNA Pos. Tick Rate” indicates the ratio of hedgehogs infested with SFTSV RNA positive ticks.

Additional *E. amurensis* hedgehog serum samples were collected from trapping studies conducted in other SFTS endemic areas, including Weifang City of Shandong Province, Linfen City of Shanxi Province, and Xinyang City of Henan Province. SFTSV antibodies were detected in 9/35 (25.7%), 2/6 (33.3%) and 2/8 (25%) of hedgehogs from Weifang City, Linfen City and Xinyang City, respectively; 11.1%, 16.7% and 0 of hedgehogs were tested positive for SFTSV RNA, respectively; and 11.1%, 50% and 12.5% of hedgehog were infected by ticks positive for SFTSV RNA (Fig. 1E and 1F). We believe these results strongly support our hypothesis that hedgehogs play an important role in the natural circulation of SFTSV.

## The susceptibility of hedgehogs to experimental infection with SFTSV

Four male and female *E. amurensis* hedgehogs (6-12 months old) were inoculated with 4 × 10^6^ FFU of SFTSV by intraperitoneal (i.p.) route. A viremia of around 9 d was observed in all animals, with peak titers of 3.1 log10 RNA copies/μl at d 3-6, suggesting viral multiplication. Two *E. amurensis* hedgehogs showed a mild weight loss of less than 25% by 9 d (Fig. 2A-2B). Neutralizing antibody titer against SFTSV in the sera of two *E. amurensis* hedgehogs were measured at 20, 30 and 40 d post-infection (dpi). In contrast to the stable humoral immune response of experimental dogs(Niu et al., 2013), the neutralizing antibody titer decreased quickly and was almost eliminated by d 40 (Fig. 2C).

**Figure 2.**
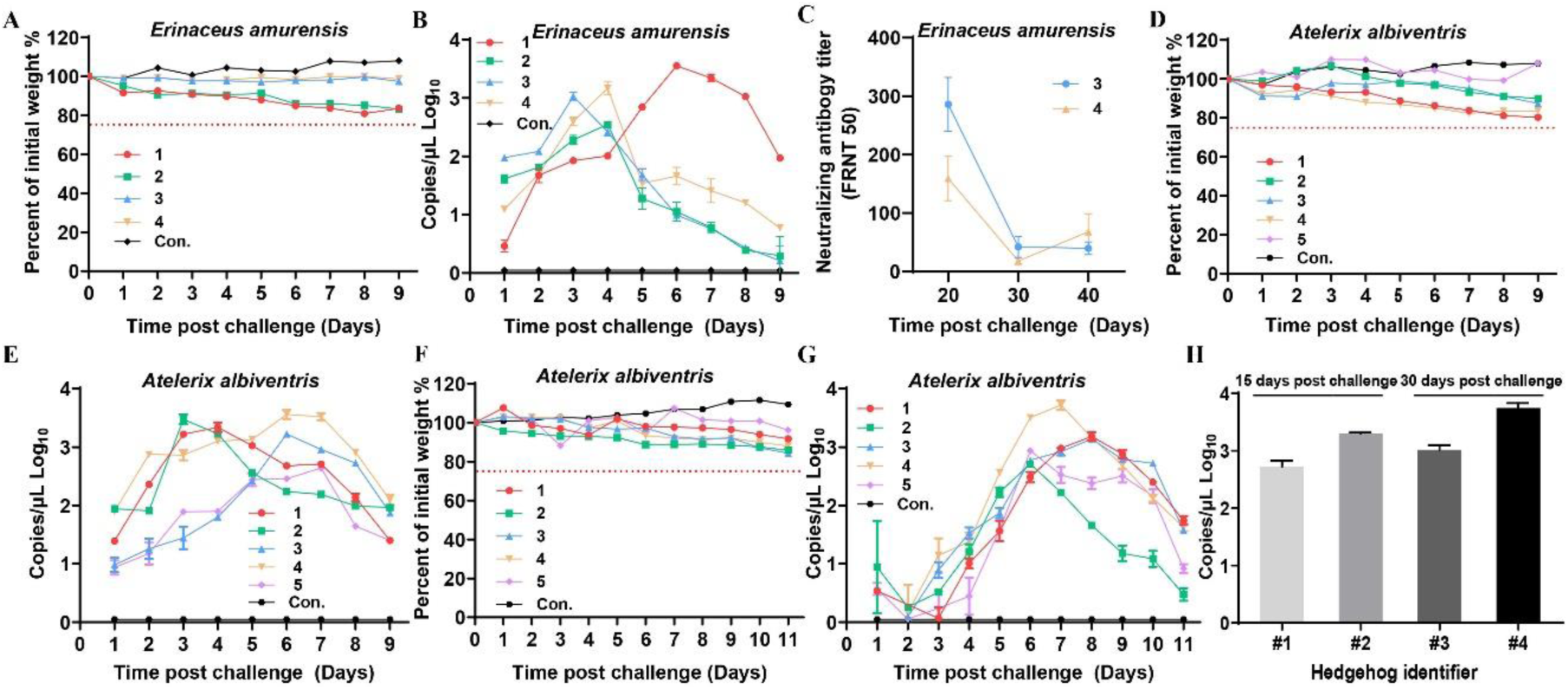
SFTSV viremia in experimentally infected *E. amurensis* and *A. albiventris* hedgehogs. Hedgehogs were challenged (i.p. or i.s.) with 4 × 10^6^ FFU of SFTSV Wuhan strain and then monitored for weight change (A and D and F) and viremia (B and E and G), tested by Real-time PCR as RNA copies per ul of serum (B and E) (error bars represent SD). Control (Con.) was mock infected with PBS. (C) SFTSV neutralizing antibody titer in two *E. amurensis* hedgehogs were monitored at 20, 30, 40 dpi. (H) Hibernation extended the course of SFTSV viremia in *A. albiventris*. Four hedgehogs were challenged (i.p.) with 4 ×10^6^ FFU of SFTSV and then kept at 4 °C to trigger hibernation. Viremia in #1 and #2 was monitored 15 dpi, while #3 and #4 at 30 dpi (error bars represent SD).

Groups of five male and female *A. albiventris* hedgehogs (6-12 months) were inoculated with 4 × 10^6^ FFU of SFTSV by intraperitoneal (i.p.) and subcutaneous (i.s.) route, respectively. A viremia of around 9 to 11 d was observed in all ten animals, with peak titers of 3.2 log10 RNA copies/μl at d 3-7 for i.p. route and titers of 3.1 log10 RNA copies/μl at d 6-8 for i.s. route, respectively (Fig. 2D-2G). Most animals showed mild wight loss of less than 20%. These results suggest that *E. amurensis* and *A. albiventris* hedgehogs could develop similar viremias independent of inoculation routes.

There was a possibility that the observed weight loss could have been iatrogenic, since the blood samples were taken directly from the heart. Heart sampling was necessary because rapid blood coagulation makes sampling from superficial veins very difficult in hedgehogs(Lewis, 1976). To test whether heart sampling was the cause of the weight loss, a further six *A. albiventris* were i.p. inoculated with SFTSV as above and bled only twice on d 7 and d 14, instead of daily. We found that only one hedgehog showed a weight loss of 20% and recovered by d 16 (Fig. S1A). In these six hedgehogs, SFTSV viremia was detected at d 7 and disappeared at d 14 (Fig. S1B). Overall, these results demonstrate that both *E. amurensis* and *A. albiventris* can develop a similar SFTSV viremia after experimental infection, without significantly compromising their overall health. However, *E. amurensis* hedgehogs are shy and easy to die during transport because of the stress response. Thus, we performed most of the following experiments with *A. albiventris* due to the stable supply in the local pet store.

### SFTSV viremia during the hibernation of hedgehogs

Four *A. albiventris* were inoculated with 4 × 10^6^ FFU of SFTSV and kept at 4°C to trigger hibernation. Two of the hedgehogs came out of hibernation at d 15 with viremias of 2.7 and 3.3 log10 RNA copies/μl respectively, whilst the other two hedgehogs continued in hibernation until d 30 with viremias of 3.0 and 3.7 log10 RNA copies/μl. All the viremias measured in these hibernating hedgehogs were comparable to the peak virus titers previously measured in the non-hibernating hedgehogs (Fig. 2H). However, the duration of viremia in these 4 hibernating hedgehogs was much longer than that recorded in the non-hibernating hedgehogs, suggesting that hibernation could potentially extend the course of SFTSV viremia in hedgehogs and contribute to the overwintering of SFTSV in the field.

### SFTSV induced pathology in hedgehogs

To assess the pathological changes in hedgehogs resulting from SFTSV infection, six *A. albiventris* were i.p. inoculated with 4 × 10^6^ FFU of SFTSV. Two animals were sacrificed at 3 days, 6 days and 2 months post-infection and their organs were removed for viral RNA evaluation and hematoxylin and eosin (H&E) staining. A robust viremia was detected on d 3 and d 6 but was negative at 2 months after infection. The highest level of viral RNA was observed in the spleen, followed by the blood, while the lowest was in the heart (Fig. 3A). H&E-stained slides from the spleen showed hemorrhagic necrosis and lymphopenia at d 3 and d 6. The severity of the lesions was assessed as +++ and ++++ on day 3 and 6 respectively but the lesions had largely recovered by 2 months with a severity score of ++ (Fig. 3B). These results further confirmed that hedgehogs show a high tolerance to SFTSV without obvious long-term or permanent pathological changes.

**Figure 3.**
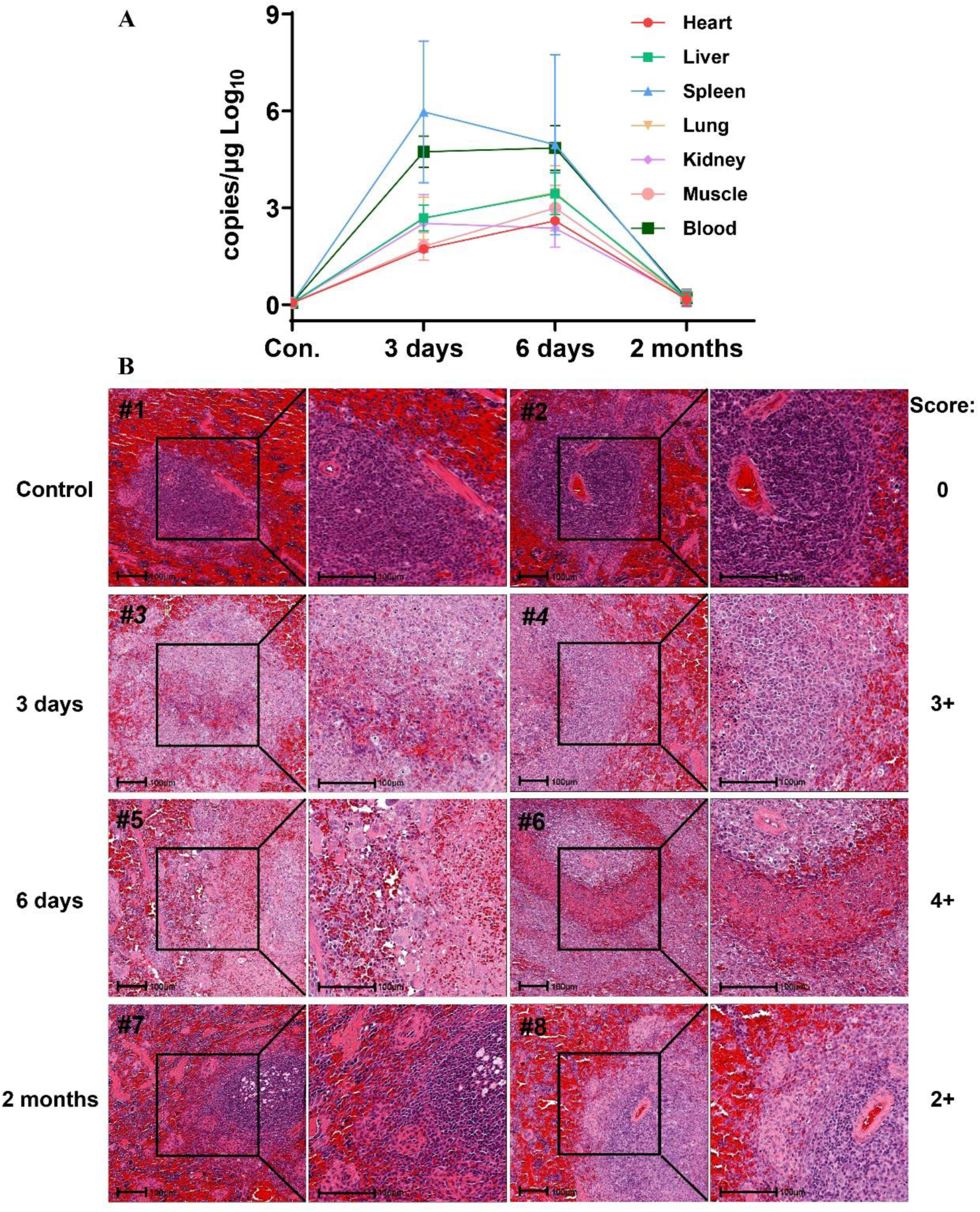
Pathology of SFTSV infected *A. albiventris* hedgehogs. Six hedgehogs were challenged (i.p.) with 4 × 10^6^ FFU of SFTSV Wuhan strain and two were mock infected by PBS as control (Con.). Two hedgehogs were killed at indicated time points to test the viral load in the organs (A) and pathology of the spleen (B). (A) SFTSV viral load in organs was measured by Real-time PCR. (B) Spleen samples were H&E stained for the pathological interpretation, with the severity of pathological changes shown beside the image. Size bars indicate 100 μm.

### Transmission of SFTSV by *H. longicornis* between ticks and hedgehogs

Lab-adapted *H. longicornis* ticks and *A. albiventris* hedgehogs were used to model the natural transmission of SFTSV hypothesized to occur in the wild. Naïve *H. longicornis* nymphs were fed on hedgehogs infected by i.p. inoculation with 4 × 10^6^ FFU of SFTSV at d 0. Viremia of 3.8 log10 RNA copies/μl was detected in hedgehogs at d 5 (Fig. 4A) and fully engorged nymphs dropped off between d 4 to 8. After molting, the adult ticks tested 100% positive for SFTSV, with a level of 7.2 log10 RNA copies/mg tick (Fig. 4B).

**Figure 4.**
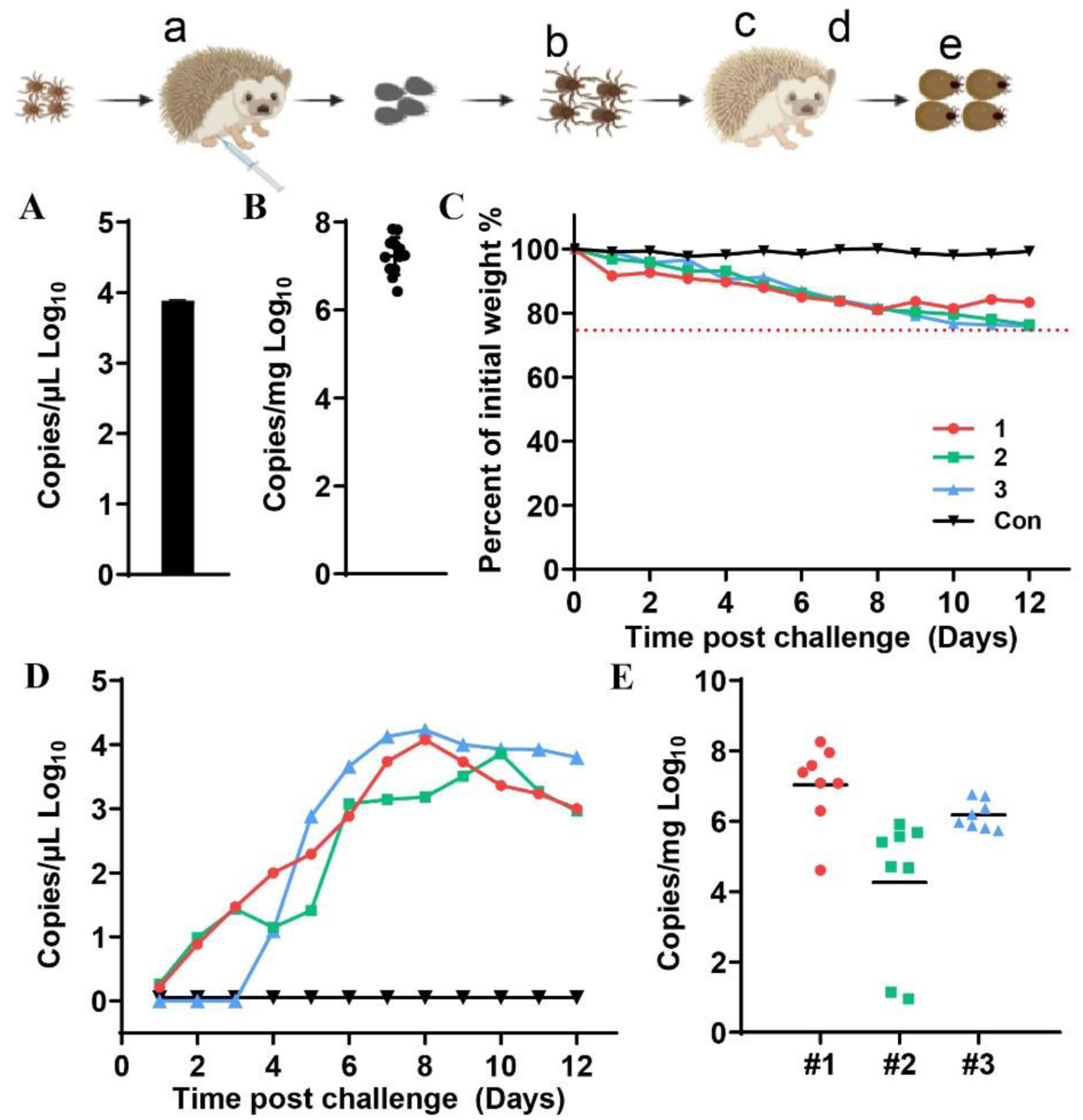
Transmission of SFTSV between *H. longicornis* ticks and *A. albiventris* hedgehogs. Hedgehogs were i.p. inoculated with 4×10^6^ FFU of SFTSV Wuhan strain and naïve nymphs were fed on the hedgehogs at the same time. (A) SFTSV viremia in the hedgehogs at 5 dpi as measured by Real-time PCR. (B) SFTSV RNA copies in the adult ticks after molting as shown by RNA copies per mg of tick. (C) Weight change and (D) SFTSV viremia in naïve hedgehogs bitten by SFTSV-carrying adult ticks were monitored for 12 d. (E) SFTSV RNA level in the engorged adult ticks from three hedgehogs as shown by RNA copies per mg of tick. Each dot indicates one tick (error bars represent SD). # indicates hedgehog identifier. The experimental process is graphically displayed above the plots and read from left to right, with the lower-case letters (a-e) corresponding to the upper-case letters of the main panels (A-E).

Three naïve hedgehogs were each fed on by eight SFTSV-carrying adult ticks at d 0. Weight and viremia were monitored for 12 d. A slow weight loss of less than 25% was observed by d 12 and the viremia peaked between d 8-10 at 4.1 log10 copies/μl. After peaking, the viraemia decreased slowly until the 3 hedgehogs were euthanized on d 12 (Fig. 4C and 4D). The fully engorged ticks were collected between d 7 to 10 and then tested. All 24 ticks were still positive for SFTSV RNA (Fig. 4E). We believe that these data strongly suggest that SFTSV can be efficiently transmitted between hedgehogs and *H. longicornis* ticks, and that transstadial transmission occurs within *H. longicornis* ticks.

### Hedgehogs are the amplifying host for SFTSV

SFTSV can be transmitted both transovarially and transstadially in *H. longicornis*, however, a decreased efficiency has been observed during passaging(Zhuang et al., 2018). Thus, an amplifying host will be necessary to improve the transmission efficiency. SFTSV-positive adult *H. longicornis* ticks were prepared as described above with 100% efficiency (Fig. 5A and 5B).

**Figure 5.**
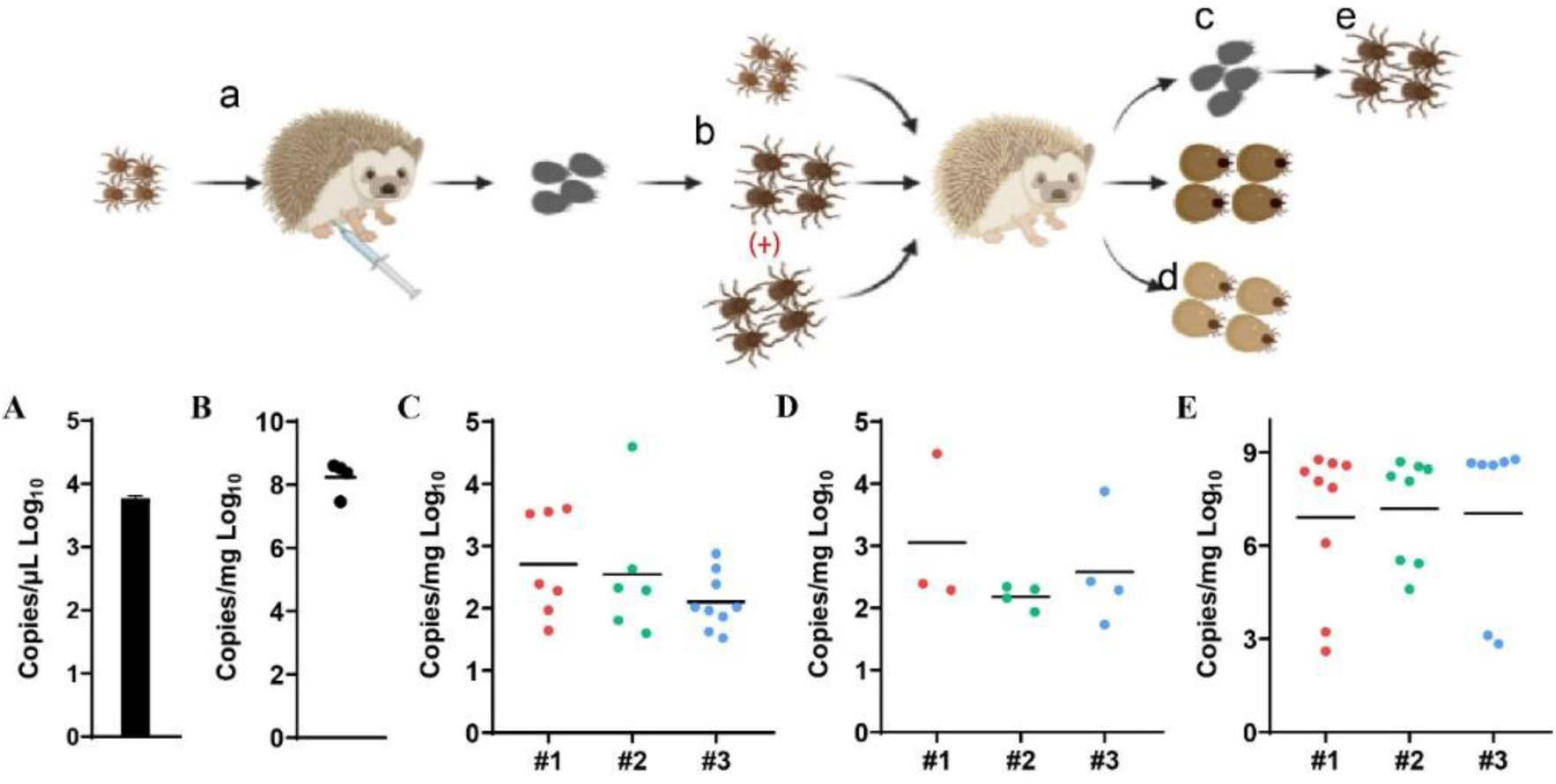
Naïve *H. longicornis* ticks infected by SFTSV through co-feeding with SFTSV positive ticks on naïve *A. albiventris* hedgehogs. Hedgehogs were i.p. inoculated with 4×10^6^ FFU of SFTSV Wuhan strain and nymph ticks were fed on the hedgehogs at the same time. (A) SFTSV viremia in hedgehogs at 5 dpi as measured by Real-time PCR. (B) SFTSV RNA copies in adult ticks after molting as shown by RNA copies per mg of tick. After molting, the SFTSV-carrying adult ticks and naïve nymph/adult *H. longicornis* ticks were fed on three naïve *A. albiventris* hedgehogs. The fully engorged ticks were collected 7 to 10 d post biting. (C-E) SFTSV RNA level was monitored in ticks as shown by RNA copies per mg of tick. Each dot indicates one tick. (C) Engorged nymph ticks. (D) Engorged adult ticks. (E) Adults molted from (C). The experimental process is graphically displayed above the plots and read from left to right, with the lower case letters (a-e) corresponding to the upper case letters of the main panels (A-E). (+) indicates SFTSV infected ticks.

Next, 5 of the SFTSV-carrying adult *H. longicornis* ticks were fed together with 14 to 16 naïve nymphs and 3 to 4 naïve adult ticks on each of three naïve *A. albiventris* hedgehogs. The fully engorged ticks were collected between 7 to 10 d post biting and tested for the viral RNA levels. The viral load in the engorged nymphs and previously naïve adults were 2.5 and 2.7 log10 RNA copies/mg tick respectively (Fig. 5C and 5D). After the nymphs molted, the adult ticks tested 100% positive for SFTSV, with a level of 6.9 log10 RNA copies/mg tick (Fig. 5E). Thus, these results suggest that hedgehogs could be acting as an amplifying host for SFTSV.

### Natural circulation of SFTSV in the urban area

The density of hedgehogs in two rural villages from Daishan County in Zhejiang Province and in two urban communities from Chaoyang and Haidian District in Beijing City were found to be very similar (Table 1). To investigate the potential natural circulation of SFTSV in the urban setting, we carried out field surveys on small mammals and parasitic ticks at two locations in Beijing City in 2021, one small park surrounded by up-market gated communities in Shunyi District and the Olympic Forest Park in Chaoyang District, where parthenogenetic *H. longicornis* ticks were discovered in a previous survey in 2019 (Fig. 6A)(X. Zhang et al., 2022). Six *E. amurensis* hedgehogs were caught in the Shunyi location and showed 2/6 (33%) SFTSV seroprevalence (Fig. 6D). *H. longicornis* ticks collected from vegetation at the same location tested positive for SFTSV RNA, clustering into lineage C4, similar to the strains of SFTSV collected in Xinyang City (Fig.6C). In contrast, no SFSTV RNA or antibody was detected in animal serum samples and parasitic ticks collected at the Chaoyang sample site. Through phylogenetic analysis of the whole mitochondrial sequences, we further found that the parthenogenetic *H. longicornis* ticks collected from the Shunyi and Chaoyang districts were closely related to those from Guangshan County, Xinyang City in Henan Province (Fig. 6B). These results imply that both the tick and SFTSV collected in Shunyi Distict might have originated from Xinyang City, one of the original SFTS hot spots. Although no local SFTS cases have been reported in Beijing yet, our results suggest that a population of hedgehogs and *H. longicornis* ticks could maintain the circulation of SFTSV in the urban ecosystem, which might result in urban SFTSV epidemics in the future.

**Figure 6.**
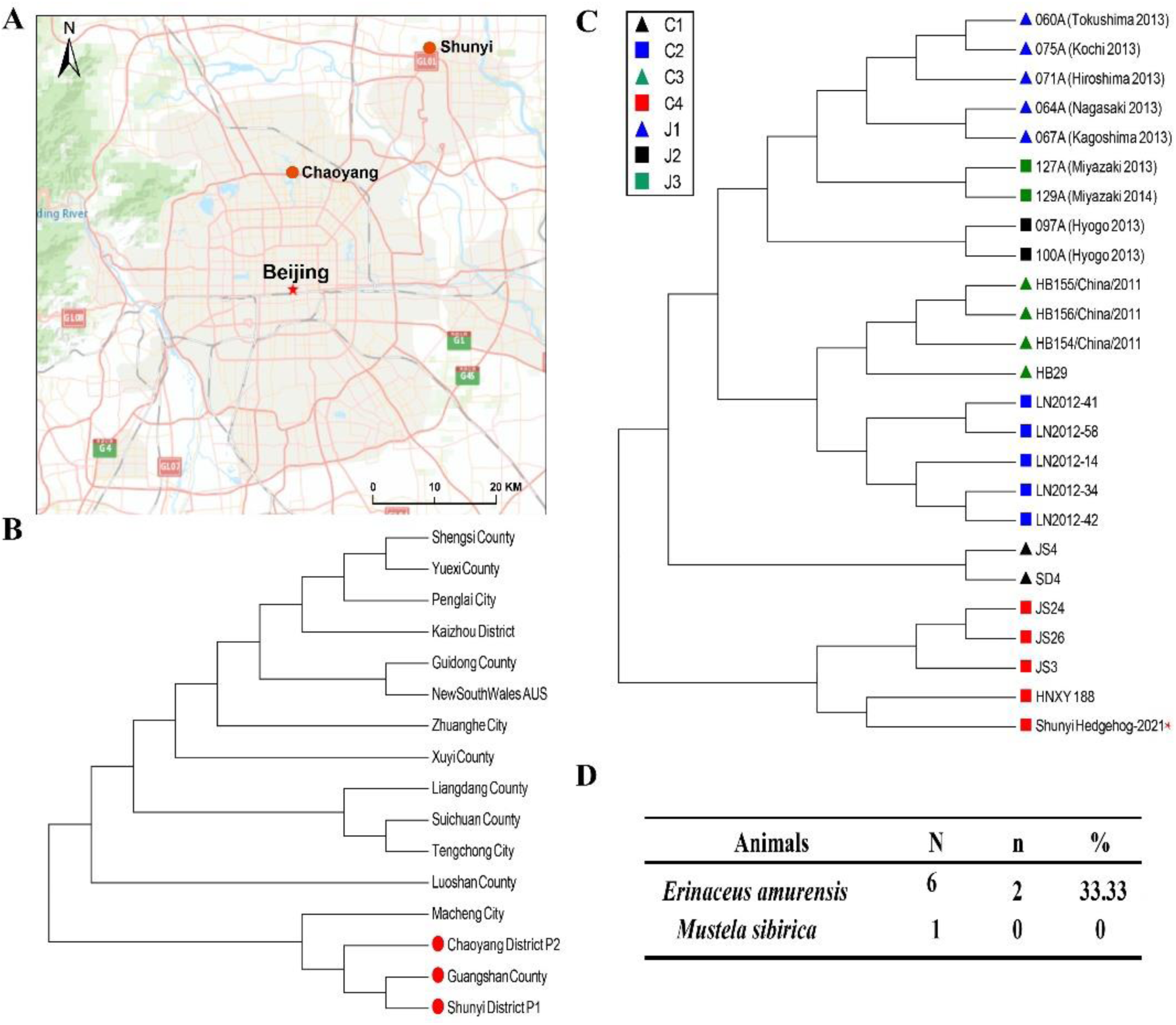
Natural circulation of SFTSV in the urban Beijing. (A) Locations studied in Shunyi District and Chaoyang District. (B) Phylogenetic analysis of the parthenogenetic population of Asian long-horned ticks. Maximum likelihood tree was established with the mitochondrial genomes of *H. longicornis* collected in Chaoyang District and Shunyi District (Chaoyang DistrictP2, accession number, OL335942 and ShunyiDistriceP1 accession number, OL335941) (red) and from SFTS endemic areas(X. Zhang et al., 2022). (C) Maximum likelihood tree was established with the L Segments of SFTSV isolate in Shunyi (Shunyi-hedgehog-2021) and isolates from SFTS endemic areas(Shi et al., 2017; Yoshikawa et al., 2015). HNXY188 was isolated from Xinyang City, Henan Province. SFTSV lineages were illustrated by colors and shapes. (D) Seroprevalence of animals caught in Shunyi district. N, number of sampled animals; n, number of sampled animals positive for SFTSV antibody; %, percentage of sampled animals positive for SFTSV antibody.

## Discussion

Viremia in the vertebrate host is important for the arbovirus to transmit from host to vector. Previous epidemiological surveys and experimental infections have revealed that many wild and domesticated animals are susceptible to SFTSV infection(Chen et al., 2019). However, these studies had similar findings that most vertebrate animals were only sub-clinically infected with SFTSV, with a limited viremia(Casel, Park, & Choi, 2021). For example, 80% of goats developed a viremia, after i.s. inoculation with 10^7^ PFU of SFTSV, which lasted for only less than 24 h(Jiao et al., 2015). Similarly, beagle dogs intramuscularly inoculated with 2.51 × 10^7^ TCID50 of SFTSV, only had a detectable viremia at d 3(S. C. Park et al., 2021). Furthermore, the efficient transmission of SFTSV between tick vectors and these potential wild animal hosts has not been proven. In this study, robust viremias of about 10^3^ RNA copies/μl were consistently detected in both native *E. amurensis* and exotic *A. albiventris* hedgehogs after i.p. or i.s. inoculation with 4 × 10^6^ FFU of SFTSV at 100% efficiency and lasted for nine to eleven days, which provides basis for the effective transmission of SFTSV from host to tick.

Hedgehogs were highly tolerated to SFTSV infection, with slight weight loss and pathology which recovered after the clearance of virus. SFTSV seroconversion was observed. In contrast to the stable humoral immune response of experimental dogs and convalescent patients(L. Hu et al., 2021; Niu et al., 2013), the antibody titers in hedgehogs decreased quickly, suggesting that studies measuring the seroprevalence of wild hedgehogs may have underestimated the true prevalence of infection and that hedgehogs might even be vulnerable to reinfection by SFTSV.

*H. longicornis* overwinters mostly as nymphs, but with an SFTSV positive rate of only 4% as measured by pool(Kim et al., 2020). Thus, we speculate that their role in overwintering of disease may be quite limited. Hedgehogs are involved in the overwintering of many pathogens during hibernation (Nosek & Grulich, 1967; Simkova, 1966) which could include SFTSV. Our results suggest that the SFTSV viremia can be extended from nine days, when non-hibernating, to at least one month during hibernation, and with a similar peak viremia to that seen in non-hibernating hedgehogs.

To meet the requirement for hedgehogs to be considered as important maintenance hosts for SFTSV, the transmission cycle between vector and host needs to be established. Using lab-adapted *H. longicornis* ticks and *A. albiventris* hedgehogs, this study conclusively showed efficient infection transmission from nymph or adult ticks to hedgehogs, efficient infection transmission from hedgehogs to nymph or adult ticks and transstadial infection transmission from nymph to adult tick. It is important to emphasize that these results were observed in 100% of tested subjects. Naïve nymph and adult *H. longicornis* ticks co-feeding with SFTSV-infected adult ticks on naïve hedgehogs were also 100% infected. We believe our results clearly show that hedgehogs fulfill the requirements to be considered the competent amplifying hosts for SFTSV. It’s still plausible that other animals or birds could also maintain the natural circulation of SFTSV. For example, experimentally inoculated spotted doves (*Streptopelia chinensis*) can develop SFTSV viremia, however, the transmission between *H. longicornis* ticks and spotted doves is not proven(Li et al., 2019).

To conclude that hedgehogs are the major amplifying hosts of SFTSV in the real world, abundance, tick-association, geographic distribution in areas of transmission, and field exposure need to be investigated. Our initial survey in SFTSV endemic Daishan Island and non-endemic Xiushan Island reveals that the existence of hedgehogs was corelated to SFTSV transmission. The epidemiological surveys we conducted in four SFTSV endemic provinces consistently showed high SFTSV seroprevalence and that the population density of hedgehogs in SFTSV endemic areas can be much higher than 60 animals per km^2^. Also, Hedgehogs are heavily infested by ticks including *H. longicornis* with a density of 145 ticks per animal as observed in Daishan Island. Hedgehogs are widely distributed across farms and rural communities, which contain the people most likely to be bitten by *H. longicornis* carrying SFTSV(Li et al., 2014; Y. Sun et al., 2017). Furthermore, hedgehogs share the same environment as domestic animals such as dogs, goats, and cows, which are also natural hosts for *H. longicornis* ticks and showing high seroprevalence to SFTVS. Thus, it is possible that humans and domestic animals get similarly infected by ticks which had previously fed on SFTSV-positive hedgehogs at an earlier stage in their life cycle. As previously stated, there are few large wild animals in SFTSV endemic areas in China and the most common animals are rodents and insectivores. Tests on rodents have shown that they are not capable of maintaining infection(Matsuno et al., 2017). Our results conclusively show that of the mammals present in rural China, the hedgehogs meet all the requirements of major amplifying hosts for SFTSV.

Beijing is the capital of China and is abundant in *H. longicornis* ticks and *E. amurensis* hedgehogs, between which local transmission of SFTSV were detected recently in an urban community in Shunyi District. Considering that there are no livestock, poultry, and stray dogs in this area, it is reasonable to infer that SFTSV circulation can be maintained by just hedgehogs and *H. longicornis* ticks in an urban area. So far, no human SFTS cases reported, but further surveillance is warranted.

SFTSV may also spread to other countries with competent hosts and vectors. *E. europaeus* hedgehogs were introduced to New Zealand by human intervention(Brockie, 1975; ISAAC, 2005). The summer density of hedgehogs in three studies in New Zealand was estimated at between 250 hedgehogs/km^2^ (Brockie, 1957; Parkes, 1975) and 800 hedgehogs/km^2^ (Campbell, 1973). In addition *H. longicornis* ticks are common in New Zealand and are all parthenogenetic (Figure S2)(Heath, 2016). New Zealand is also on the East Asian-Australian flyway. Therefore, New Zealand might be considered to have a high risk of SFTSV disease incursion, likely through SFTSV positive *H. longicornis* ticks infested in migratory birds.

In conclusion, our data strongly support our initial hypothesis that hedgehogs can maintain the natural circulation of SFTSV in rural areas. The high density and wide distribution, the high-level susceptibility and tolerance of hedgehogs to SFTSV, the heavy *H. longicornis* infestation rates and the ability to amplify the infection level of feeding ticks are all compelling evidence that hedgehogs are the major wildlife amplifying host of SFTSV. Furthermore, evidence that SFTV is already circulating in ticks and hedgehogs from one urban area of Beijing means that an urban epidemic of SFTS could happen quite soon. More research and surveillance are needed to reduce SFTSV risks in both rural and urban areas.

## Materials and Methods

### Ethics statement

All animal studies were carried out in strict accordance with the recommendations in the Guide for the Care and Use of Laboratory Animals of the Ministry of Science and Technology of the People’s Republic of China. The protocols for animal studies were approved by the Committee on the Ethics of Animal Experiments of the Institute of Zoology, Chinese Academy of Sciences (Approval number: IOZ20180058).

### Animal trapping and sample collection

Animal sampling took place in Daishan City (121°30′-123°25′E, 29°32′ ∼ 31°04′N), Zhejiang Province, Weifang City (118°10’-120°01’E, 35°32’-37°26’N), Shandong Province, Xinyang City (113°45″-115°55″E, 30°23″-32°27″ N), Henan Province, Linfen City (110°22′-112°34′E, 35°23′-36°57′N), Shanxi Province, and Beijing City (39°26’-41°03’E, 115°25’-117°30’N), China. The animals were captured using rodent capture cages (cage size: 14 × 14 × 26 cm) baited with fried bread sticks for three nights at each site (trappings varied between 30 and 50 traps/night depending on the availability of sites in the area). Cages were deposited into fields and collected the next morning (FERNANDO TORRES-PÉREZ, 2004). Animals were anesthetized by inhalation using Isoflurane with a dose of 1 mL per kilogram weight in a closed container. Blood samples were drawn from heart, and animals were released after blood collection. Blood samples were centrifuged at 3000 g for 10 minutes and the serum was transferred to small vials, which were kept at -80°C until analysis.

### Virus and cells

SFTSV Wuhan strain (GenBank accession numbers: S, KU361341.1; M, KU361342.1; L, KU361343.1) and rabbit anti-SFTSV-NP polyclonal antibody were provided by Wuhan Institute of Virology, Chinese Academy of Sciences. Vero cells (African green monkey kidney epithelial cells) were obtained from American Type Culture Collection (ATCC) and maintained in Dulbecco’s modified Eagle’s medium (DMEM, Hyclone, US) supplemented with 8% FBS and penicillin (100 U mL^-1^), streptomycin (100 μg mL^-1^; Gibco) and L-glutamine in a 37°C incubator supplemented with 5% CO_2_. SFTSV was propagated at 37°C in Vero cells at a multiplicity of infection of 0.1 and cultivated for 4 d. Cell culture supernatant was collected at 4 dpi and stored at -80°C as the working virus stock for animal studies.

### Virus titration

Focus-forming assay was performed in Vero cells to titrate the viral titers. Cells were seeded in 96-well plates at 10^4^ cells/well in triplicates 24 h before infection. The virus samples were diluted 10-fold in DMEM with 2% FBS. After the removal of culture media, a diluted viral solution was added to the cells. Three hours later, the cells were washed once and incubated with DMEM plus 2% FBS and 20mM NH_4_Cl at 37°C. 2 d post-infection, the cells were fixed with cold methanol and stained using a rabbit anti-SFTSV-NP polyclonal antibody at 1:700 dilution and Alexa 488-labeled goat anti-rabbit IgG at 1:700 dilution. Viral titers were examined under a fluorescent microscope and calculated by Reed–Muench method.

### ELISA for SFTSV antibody detection

Serum samples from animals were tested for SFTSV antibodies including IgG and IgM with a commercial double antigen sandwich ELISA kit from Nanjing Immune-detect Bio-tech Co., Ltd. (Jiangsu, China).

### Experimental Infection

All experimental infection study was conducted in a Bio-safety Level-II Animal Laboratory in the Beijing Institute of Microbiology and Epidemiology, Academy of Military Medical Sciences. Six to twelve months old male and female (1:1) African pygmy hedgehogs were purchased from Longchong Pet in Beijing. Six to twelve months old male and female (1:1) Amur hedgehogs were purchased from Heze animal store in Shandong Province. All animals were tested for SFTSV seroprevalence by ELISA before experiments (Figure S3). Following acclimation, hedgehogs were challenged with 4× 10^6^ FFU of SFTSV Wuhan strain via i.p. injection or i.s. injection, with the 200 ul volume divided between two injection sites. Bodyweight and clinical symptoms were monitored. Hedgehogs were assigned a clinical score of increasing severity: 1, unfeeding; 2, hunched posture; 3. green faeces;4, moribund. Hedgehogs with a score of 3 or a weight loss of more than 25% were humanely euthanized.

For the virus transmission studies between host and vector, naïve ticks were fed on African pygmy hedgehogs which were i.p. inoculated with 4 × 10^6^ FFU of SFTSV Wuhan strain at d 0, and the ticks collected when they naturally detached. And for the transmission studies between vector and host naive African pygmy hedgehogs were bitten by infected ticks. Bodyweight and clinical symptoms in bitten hedgehogs were monitored.

For the hibernation experiment, Hedgehogs were challenged with 4 × 10^6^ FFU of SFTSV Wuhan strain and then spent half month or one month in 4°C refrigerator. Two of the hedgehogs were waken up at d 15 and another two were waken up at d 30. The blood samples were taken from the heart for viral RNA detection.

### SFTSV-RNA extraction and real-time RT-PCR

Total RNA prepared from the homogenates of the ticks and the blood samples collected from hedgehogs’ heart were extracted using TRIzol reagent (Thermo Fisher Scientific, USA) or the RNeasy kit (Qiagen, Germany) according to the manufacturer’s instructions. Samples were analyzed using a One-Step SYBR PrimerScript reverse transcription (RT)-PCR kit (TaKaRa, Japan) on Applied Biosystems QuantStudio. Each sample was measured by triplicate. The primers were designed as previously described(Dong et al., 2019). Conditions for the reaction were as follows: 42°C for 5 min, 95°C for 10 sec, 40 cycles at 95°C for 5 sec, and 60°C for 20 sec.

### Pathological lesions in SFTSV-Infected hedgehogs

Six African pygmy hedgehogs were inoculated with 4 × 10^6^ FFU of the Wuhan strain of SFTSV by i.p. route and two were killed at each time point of d 3, 6 and months 2 post infection for analysis of viremia and pathology. Two mock-infected control hedgehogs were killed at d 0. For histopathological evaluation, spleens were rapidly removed, fixed in 4% PFA at room temperature for 7 d, and routinely processed for paraffin embedding. Coronal, 4–5 μm thick serial sections were performed and selected sections were stained with H&E for light microscopy examination. Images were obtained using a Nikon Eclipse 50i Light Microscope (Nikon, Tokyo, Japan) or Olympus BX60 microscope (Shinjuku, Tokyo, Japan)

### Identification of tick species and phylogenetic analysis

Ticks were identified based on morphological characteristics, visualized through a light microscope, with further molecular confirmation in the laboratory by sequencing the mitochondrial 16S ribosomal RNA (16S rRNA) gene. The primers were as follows: (16S-1) CTGCTCAATGATTTTTTAAATTGCTGTGG (Forward primer) and (16S-2) CGCTGTTATCCCTAGAGTATT (Reverse primer). A single leg was removed from each tick for the molecular analysis to confirm identification. Phylogenetic analysis was performed using the whole mitochondrial genomes. Tick DNA was extracted using the MightyPrep reagent for DNA Kit (Takara, Japan) according to the manufacturer’s instructions. The mitochondrial DNA were sequenced by next generation sequencing (Tsingke Biotech, Beijing, China) and deposited in GenBank (Parthenogenetic *H. longicornis* in Shunyi District: OL335941; Chaoyang District: OL335942). The phylogenetic tree was constructed using the maximum likelihood method, MEGA-X with the bootstrap value set at 1000.

### SFTSV sequencing and phylogenetics analysis

The SFTSV sequences from Shunyi *H. longicornis* ticks were amplified using (L4010-F:GGAACTCTCAGCCACTCTGTT; L4963-R:GTAGAGAAGGCCTCTATGATC) and sequenced by Tsingke Biotech, Beijing, China, and then deposited in the GenBank (Accession number: OL518989). SFTSV sequences in the phylogenetics analysis were downloaded from GenBank(Shi et al., 2017; Yoshikawa et al., 2015). The phylogenetic trees were constructed using the maximum likelihood method, MEGA program. The confidence of the tree was tested using 1000 bootstrap replications.

## Acknowledgments

We gratefully thank the following funders: the State Key Research Development Program of China (2019YFC12005004, 2019YFC12005001), the Strategic Priority Research Program of Chinese Academy of Sciences (Grant No. XDPB16), the key program of Chinese Academy of Sciences (CAS) (KJZD-SW-L11), Natural Science Foundation of Zhejiang Province (NO. LY21H100002), and the Open Research Fund Program of State Key Laboratory of Integrated Pest Management (IPM2109). We also thank Rachel Summers, Massey University for constructing the New Zealand hedgehog distribution map.

## Supplementary Information for

**Figure S1.**
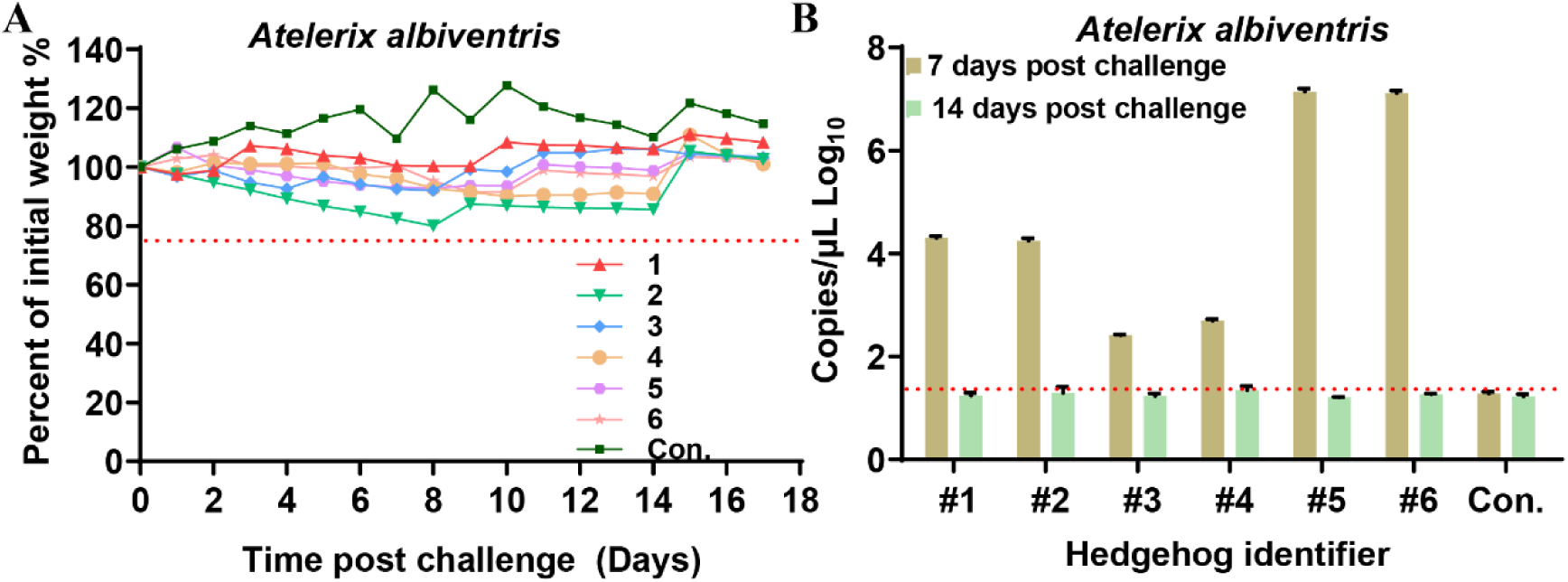
SFTSV viremia and weight change in experimentally infected *A. albiventris* hedgehogs bled twice. Hedgehogs were challenged (i.p.) with 4 × 10^6^ FFU of SFTSV Wuhan strain and then monitored for weight loss everyday (A) and viremia on d 7 and 14, measured by Real-time PCR as RNA copies per ul of serum (B) (error bars represent SD). Each line and bar indicate one hedgehog.

**Figure S2.**
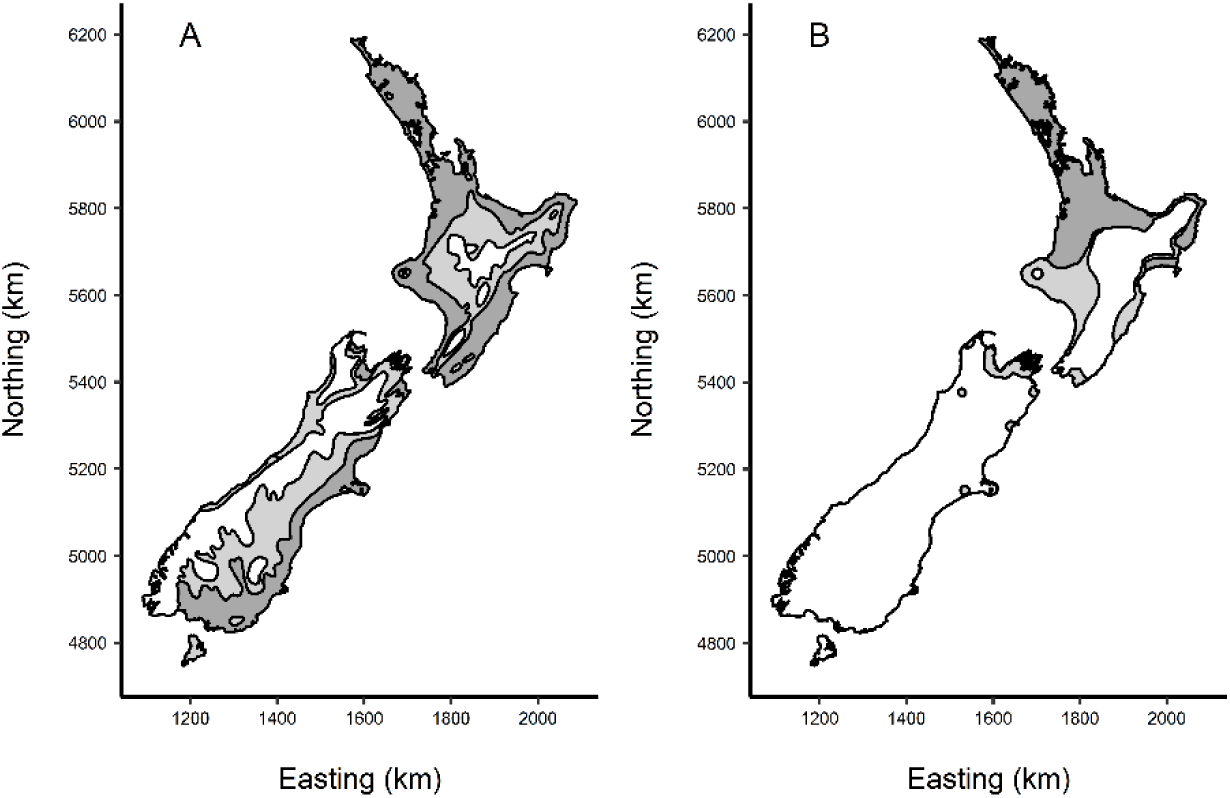
(A) The distribution and relative abundance of hedgehogs (*Erinaceus europaeus L*.) in New Zealand modified from (Brockie, 1975). In the dark grey areas hedgehogs are numerous, in the light grey areas they are few and in the white areas they are rare or absent. (B) The distribution of *Haemaphysalis longicornis* in New Zealand modified from (Heath, 2016). The dark grey areas are high risk, the light grey areas are low risk, and the white areas are zero risk of *H. longicornis* Infestation. (Reproduced with permission of Elsevier and Copyright Clearance Center).

**Fig. S3.**
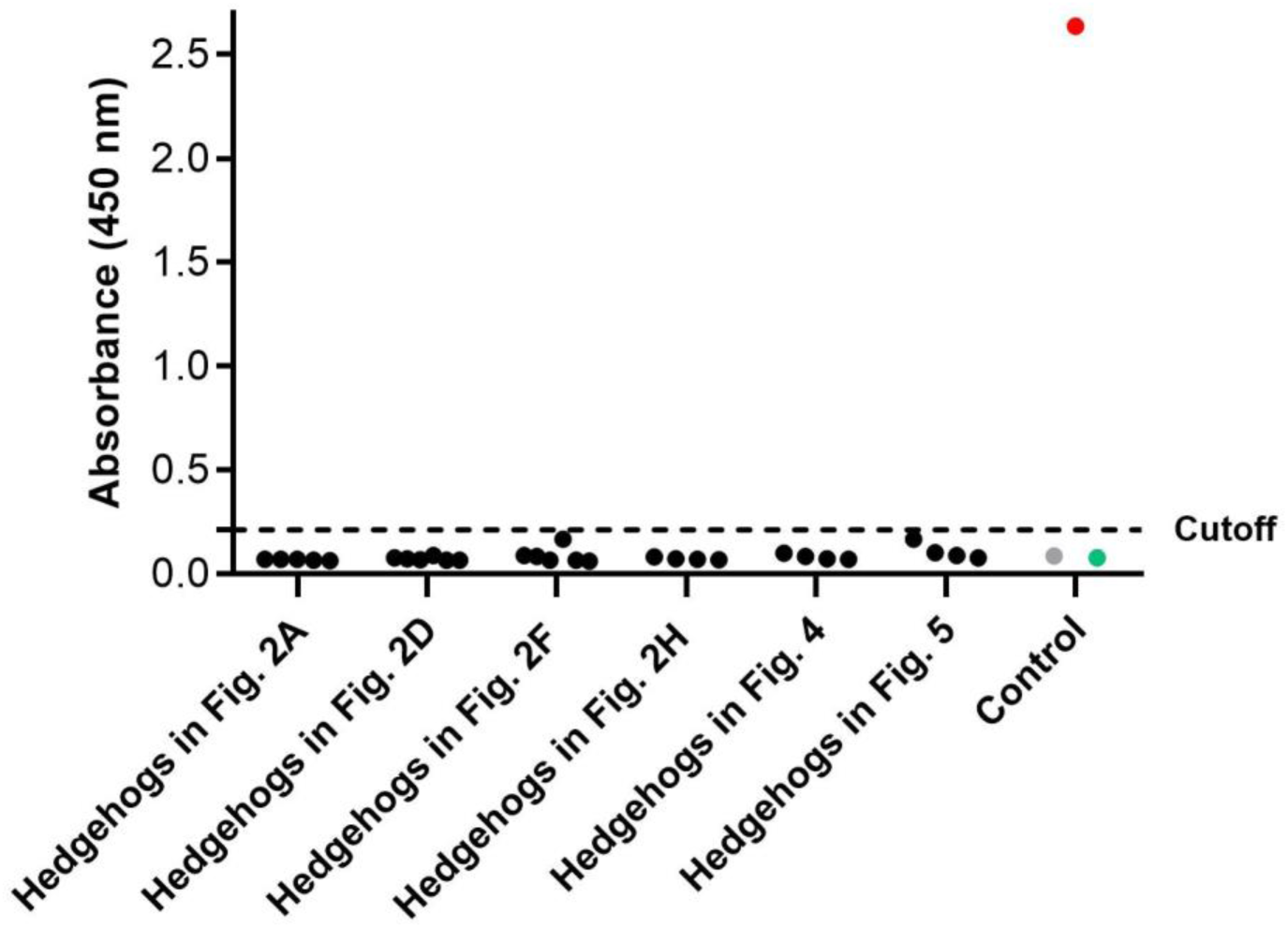
Seroprevalence of SFTSV in all hedgehogs prior to experimental infection. The serum samples were tested for SFTSV IgG and IgM antibodies prior to experimental infection with a commercial double antigen sandwich ELISA kit. Cutoff = 0.748×Negative OD +0.146. Red dot indicates positive sample, green dot indicates negative sample and gray dot indicates blank control.

